# Machine Learning to Identify Molecular Markers for Metabolic Disease Development Using Mouse Models

**DOI:** 10.1101/2023.03.11.532149

**Authors:** Guiyan Yang, Rex Liu, Shahbaz Rezaei, Xin Liu, Yu-Jui Yvonne Wan

**Affiliations:** Department of Medical Pathology and Laboratory Medicine, University of California, Davis, Sacramento, CA, USA; Department of Computer Science, University of California, Davis, CA, USA

**Keywords:** Diet, Aging, FXR, Metabolic liver disease, Bile acid, Noninvasive biomarker, Predictor, Omics

## Abstract

**Background:** Aging, Western diet (WD) intake, and bile acid (BA) receptor farnesoid X receptor (FXR) inactivation are risk factors for metabolic disease development including nonalcoholic fatty liver disease (NAFLD) and chronic inflammation-related health issues such as dementia. The progression of the metabolic disease can be escalated when those risks are combined. Inactivation of FXR is cancer prone in both humans and mice. The current study used omics data generated within the gut-liver axis to classify those risks using bioinformatics and machine learning approaches.

**Methods:** Different ages (5, 10, and 15 months) of wild-type (WT) and FXR knockout (KO) male mice were fed with either a healthy control diet (CD) or a WD since weaning. Hepatic transcripts, liver, serum, and urine metabolites, hepatic bile acids (BAs), as well as gut microbiota were used for risk prediction. A linear support vector machine with *K*-fold cross-validation was used for classification and feature selection.

**Results:** Increased urine sucrose alone achieved 91% accuracy in predicting WD intake. Hepatic lithocholic acid (LCA) and serum pyruvate had 100% and 95% accuracy, respectively to classify age. Association analyses showed hepatic LCA was positively associated with serum concentrations of acetone, a ketone body, and 1,3-dihydroxyacetone (DHA), but negatively correlated with serum pyruvate. Urine metabolites (decreased creatinine and taurine as well as increased succinate) or gut microbiota (increased *Dorea, Dehalobacterium*, and *Oscillospira*) could predict FXR functional status with greater than 90% accuracy. Integrated pathway analyses revealed that the predictors for diet and FXR expression were implicated in the central carbon metabolism in cancer. To assess the translational relevance, mouse hepatic transcripts were crosschecked with human NAFLD and hepatocellular carcinoma (HCC) datasets. WD-affected hepatic *Cyp39a1* and *Gramd1b* expression were associated with human HCC and NAFLD, respectively. The metabolites and diseases interaction analyses uncovered that the identified features are implicated in human metabolic diseases, mental disorders, and cancer.

**Conclusion:** The risk prediction using mouse models contributes to the identification of noninvasive biomarkers for early diagnosis of metabolic disease development.

## Background

The incidence of metabolic diseases such as metabolic syndrome and nonalcoholic steatohepatitis (NASH) is rising due to obesity. Early diagnosis is needed especially when patients are asymptomatic. Aging, Western diet (WD) intake, and bile acid receptor farnesoid X receptor (FXR) deactivation are risks for metabolic disease development [1–3]. FXR is one of the most characterized receptors for bile acids (BAs), which play pivotal roles in regulating lipid and glucose metabolism [4]. Lack of FXR induces hepatic steatosis, NASH, and liver cancer spontaneously as mice age [2]. Additionally, by deactivating the transcriptional activity of FXR, both aging and WD intake induce the development of metabolic disorders and chronic inflammation [4, 5]. Furthermore, when these risk factors are combined, the development of NASH and liver carcinogenesis is facilitated [1, 2, 6]. Similarly, patients with liver steatosis, cirrhosis, or hepatocellular carcinoma (HCC) have reduced expression of FXR [7].

Emerging evidence revealed the significance of dysbiosis in contributing to the development of metabolic liver disease via the gut-liver axis. Because BAs are generated by the metabolism of hepatic cholesterol using hepatic and bacterial enzymes, dysbiosis is always accompanied by dysregulated BA synthesis [1, 2, 6]. Additionally, via the diet-gut-brain axis, dysbiosis and dysregulated BA synthesis also have a negative impact on neuroplasticity [8–12]. Further, the gut microbiota community structure, the profile of BA, as well as the incidence of metabolic liver disease are sex different. Thus, it is important to discover microbes and metabolites within the gut-liver axis as biomarkers to predict the risks for increased metabolic burden such as aging and WD intake.

Computational modeling of multi-omics data can improve the depth and accuracy of omics analysis [13]. In this study, we used datasets including hepatic transcriptome and metabolomes from multiple sources including the liver, serum, and urine, as well as hepatic BAs, and gut microbiota, to identify predictors for risks of metabolic liver disease development. Our data showed different metabolites from different specimens have a specific power to predict dietary intake or age. Interestingly, the gut microbiota had the best predicting power to classify FXR expression status. Together, the uncovered molecular signatures within the gut-liver axis have the potential for detecting metabolic status.

## Methods

### Mouse models and data collection

Specific pathogen-free male wild-type (WT) and FXR KO mice [14] were fed on either a healthy control diet (CD TD.140415; 5.2% fat, 12% sucrose, and 0.01% cholesterol, w/w) or a Western diet (WD, TD.140414; 21.2% fat, 34% sucrose, and 0.2% cholesterol, w/w, Harlan Teklad, Madison, WI) since weaning and euthanized at the age of 5, 10, and 15 months. Mice were housed 3-4 per cage in a temperature (24°C) and light-controlled (12-hour light on and off cycle) facility. Multi-omics data were derived from mice with characterized phenotypes [1, 2, 6, 15, 16]. The sample size and experimental groups are summarized in Table S1 (Additional file 1). Experiments were performed in accordance with the NIH Guide for the Care and Use of Laboratory Animals under protocols approved by the Institutional Animal Care and Use Committee of the University of California, Davis.

### Analysis of liver, serum, and urine metabolites

Hepatic metabolites were measured by gas chromatography-time-of-flight mass spectrometry (GC-TOF-MS) using a sample size of 6 per group by the West Coast Metabolomics Center at the University of California, Davis. Serum and urine metabolites were quantified by NMR using published methods, and their concentrations were normalized by log transformation to reduce batch effects [17].

### RNA sequencing and data processing

Hepatic bulk RNA was used for library preparation followed by sequencing, which was conducted by Novogene Co., LTD (Sacramento, CA). Raw FASTQ data were mapped to GRCm39 (GENCODE release M27) and quantified using Salmon (version 1.4.0) [16]. DESeq2 (version 1.18) with the lfcShrink function in R 4.03 was used to identify differentially expressed genes (DEGs) [18].

### Microbiota data analysis

Cecal DNA was used for 16S rRNA sequencing using published methods [1]. The 16S rRNA gene (V4 region) was amplified and sequenced using Illumina MiSeq. Sequence reads were analyzed by QIIME [1, 2].

### Bile acid quantification

Hepatic BAs were quantified using a ProminenceTM UFLC system (Shimadzu) coupled to an API 4000 QTRAPTM mass spectrometer (AB Sciex) operated in a negative ionization mode based on published methods [1].

### Human databases

RNA sequencing data of 371 human HCCs and 50 normal livers were obtained from The Cancer Genome Atlas (TCGA) database. Transcriptomic data from non-alcoholic fatty liver (NAFL) and NASH, with or without fibrosis, cohorts were from the Gene Expression Omnibus (GEO) database (GSE 135251).

### Transcriptomic feature selection

Because the number of hepatic transcripts was much bigger than the sample size, feature selection was conducted to reduce noise in the dataset and speed up the training process [19]. Features were selected based on differences between groups with statistical significance (*p* < 0.05) and fold-change greater or equal to 2. To study dietary effects, 42 transcripts that commonly changed their expression levels in all 3 age groups (5, 10, and 15 months) were selected. Irrespective of diets, 256 transcripts differentially found between 5- and 15-month-old mice were selected. In addition, 105 transcripts differentially expressed in the livers of FXR KO and WT mice, irrespective of dietary and age differences, were included.

### Machine learning pipeline

Support Vector Machine (SVM), a trusted classical machine learning algorithm, is widely used in healthcare applications for its exceptional generalization capabilities [20–24]. In the initial analysis, linear SVM demonstrated high prediction accuracy compared with other machine learning algorithms such as non-linear alternatives. In addition, linear SVM is particularly suited for small sample size data analysis [19]. Thus, linear SVM was used for the risk prediction model.

To determine the importance of each feature in a linear SVM classifier, the associated coefficient was calculated, which corresponds to the orthogonal vector coordinate of the hyperplane [25]. The value of the coefficient is indicative of the feature’s effect on the final prediction of the model [26]. The absolute value of the coefficient was computed using linear SVM classifier via K-fold cross-validation (16-fold cross-validation for transcriptomic data and 20-fold cross-validation for other omics data). All features were sorted based on their importance and reported the least number of top-ranked features that yielded a prediction accuracy of 90% or higher for diet, age, or FXR expression status by evaluating the accuracy of various feature combinations.

### Pathway and network analysis

Kyoto Encyclopedia of Genes and Genomes (KEGG) pathway analysis for metabolites and transcripts was performed using MetaboAnalyst 5.0. The metabolite-disease interaction network in MetaboAnalyst 5.0 was used to explore disease-related metabolites based on Human Metabolome Database.

### Association analysis

Spearman’s correlation was used to assess the relationship between the predictors of each risk factor in this study. A significant correlation was defined when adjusted \**p* < 0.05 and \*\**p* < 0.01 using Hochberg.

### Statistics

The altered metabolites and bacteria between groups were considered by *p* < 0.05 using unpaired *t-*tests. DEGs were generated using DESeq2. Bar graphs were generated by GraphPad Prism Version 8.0 (San Diego, CA, USA). *p* values: \**p* < 0.05; \*\**p* < 0.01; \*\*\**p* < 0.001.

## Results

### Predictors of differential diet intakes

WD consumption induces NAFLD and increases body weight during the experimental time frames (5, 10, 15 months) [15]. The smallest feature numbers, which could generate the highest prediction accuracy are summarized in Table 1. Nine hepatic transcripts (*Cyp39a1, Pde5a, Csad, Gramd1b, Slc39a4, Hamp2, Loxl4, Rec8*, and *Adam11*) classified differential dietary intake with a 100% accuracy (Fig. 1A). Specifically, downregulated *Cyp39a1* together with upregulated *Pde5a, Csad*, and *Gramd1b* had 96.9% accuracy to predict WD intake (Fig. 1A).

**Fig. 1.**
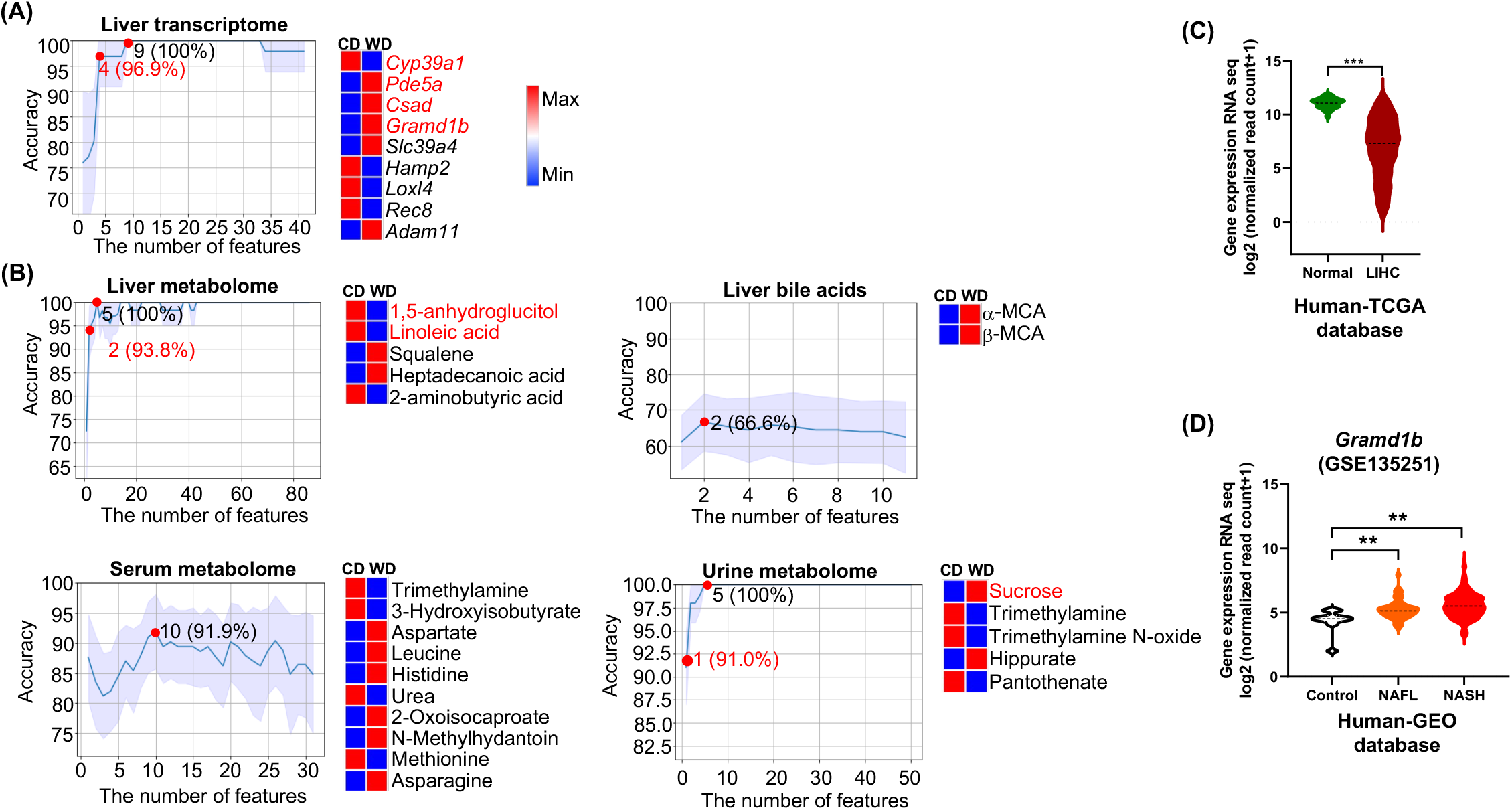
Predictors of differential diet intake based on multi-omics data. Machine learning model generated line charts on the number of features and indicated accuracy using the *K*-fold cross-validation method for (A) liver transcriptome and (B) metabolomes from the liver, serum, and urine as well as hepatic bile acids. The number of features with predictive accuracy higher than 90% and/or the number of least features that has the highest predicting accuracy is highlighted (red dot). The differences in the relative abundance of predictors between CD and WD groups are shown in heatmaps (blue and red indicate low and high levels, respectively). The order of features in the heatmap is based on the feature’s importance (coefficient value) after feature selection. (C) Human HCC patients (n = 371) have reduced *Cyp39a1* transcript compared with normal livers (n = 50) from the TCGA database. (D) A violin plot shows human NAFL (n = 51) and NASH (n = 155) cohorts have higher *Gramd1b* mRNA levels than the controls (n = 10) from the GEO database (GSE 135251). Data are expressed as the mean ± SD. \*\**p* < 0.01, \*\*\**p* < 0.001.

**Table 1.** Predictive capabilities of multi-omics in Western diet intake, aging, and FXR inactivation. Multi-omics data analyses were conducted in mice of different ages, diets, and genotypes. The least number of features that has the best predictive performance is shown for each risk factor prediction.

| Risk prediction | Western diet intake |  | Aging |  | Bile acid Receptor Inactivation |  |
| --- | --- | --- | --- | --- | --- | --- |
|  | Features | Accuracy | Features | Accuracy | Features | Accuracy |
| **Hepatic Transcripts** | 9 (4) | 100% (96.9%) | 14 (2) | 100% (90.6%) | 2 | 100% |
| **Metabolites** |  |  |  |  |  |  |
| Bile acids (liver) | 2 | 66.6% | 1 | 100% | 10 | 71.3% |
| Liver | 5 (2) | 100% (93.8%) | 20 (12) | 100% (91.7%) | 10 (1) | 100% (95.8%) |
| Serum | 10 | 91.9% | 3 (1) | 100% (95.0%) | 15 (12) | 94.5% (91.3%) |
| Urine | 5 (1) | 100% (91.0%) | 7 (3) | 100% (90.0%) | 9 (3) | 100% (95.4%) |
| **Microbiota** |  |  |  |  |  |  |
| Phylum level | 8 | 61.9% | 4 | 70.0% | 6 | 90.2% |
| Class level | 9 | 62.6% | 9 | 82.8% | 3 | 96.9% |
| Order level | 26 | 62.5% | 13 | 82.8% | 3 | 96.9% |
| Family level | 10 | 76.8% | 7 | 80.4% | 8 (3) | 98.8% (91.2%) |
| Genus level | 6 | 68.8% | 7 | 82.0% | 7 (3) | 96.2% (92.7%) |

The findings were compared with human TCGA (HCC) and GEO (NAFL, NASH) databases (GSE 135251). In consistency, the expression of *Cyp39a1*, which is involved in cholesterol clearance through BA synthesis, was downregulated in human HCC compared with normal livers (*p* < 0.001) (Fig. 1C). Additionally, *Gramd1b*, a cholesterol transporter, was consistently elevated in NAFL and NASH patients compared with healthy controls (*p* < 0.01) (Fig. 1D).

In the livers, 5 metabolites (1,5-anhydroglucitol, linoleic acid, 2-aminobutyric acid, squalene, and heptadecanoic acid) had 100% accuracy in predicting differential diet intake (Fig. 1B). Decreased 1,5-anhydroglucitol and linoleic acid yielded 93.8% accuracy in classifying diets. Hepatic a-MCA and β-MCA had 66.6% accuracy in distinguishing differential diets (Fig. 1B). Moreover, increased aspartate, leucine, histidine, 2-oxoisocaproate, N-methylhydantoin, and asparagine but decreased trimethylamine, 3-hydroxyisobutyrate, urea, and methionine found in the serum yielded 91.9% accuracy to predict WD intake. For urine metabolites, sucrose, trimethylamine, trimethylamine N-oxide, hippurate, and pantothenate achieved 100% accuracy in predicting diet. Increased urine sucrose alone had 91% accuracy in predicting WD intake (Fig. 1B).

Integrated pathway analysis uncovered that serum leucine, methionine, histidine, asparagine, and aspartate were involved in the central carbon metabolism in cancer and aminoacyl-tRNA biosynthesis (Fig. 1SA). Serum aspartate and histidine as well as urine pantothenate were involved in β-alanine metabolism (Fig. 1SA).

Network analysis showed that reduced hepatic 1,5-anhydroglucitol and linoleic acid were related to Alzheimer’s disease (Fig. 1SB). Increased urine sucrose was related to lung cancer. Increased urine TMAO was associated with many diseases including schizophrenia, propionic acidemia, maple syrup urine disease, lung cancer, and dimethylglycine dehydrogenase deficiency (Fig. 1SB).

Spearman’s correlation analysis revealed that urine sucrose was negatively associated with hepatic 1,5-anhydroglucitol and linoleic acid. Interestingly, increased urine sucrose was also negatively associated with the expression levels of *Cyp39a1*, but positively correlated with *Pde5a, Gramd1b*, and *Csad.* The decreased serum 3-hydroxyisobutyrate was positively associated with hepatic linoleic acid but negatively correlated with the expression level of *Gramd1b* (Fig. 2S). The key functions or the known roles of those transcripts and metabolites are summarized in Table S2 (Additional file 3) and Table S3 (Additional file 4), respectively.

### Age classification

Under the influence of an unhealthy diet, aging further reduces metabolic efficiency. Thus, there was a temporal effect of WD intake, and 15-month-old WD-fed mice had the most severe NAFLD [6]. The machine learning model revealed that downregulated hepatic *Zbtb16* and upregulated *Rps27rt, Naip2, Cyp46a1, Mmd2, AA792892, A4gnt, Cdh19, Pclo, Zfp677, Cyp3a11, Hsf2bp, Kcnj16, Mfsd2a*, yielded 100% accuracy to differentiate 15 and 5 months of ages (Fig. 2A). Moreover, two transcripts *Zbtb16* and *Rps27rt* had 90.6% accuracy in classifying age.

**Fig. 2.**
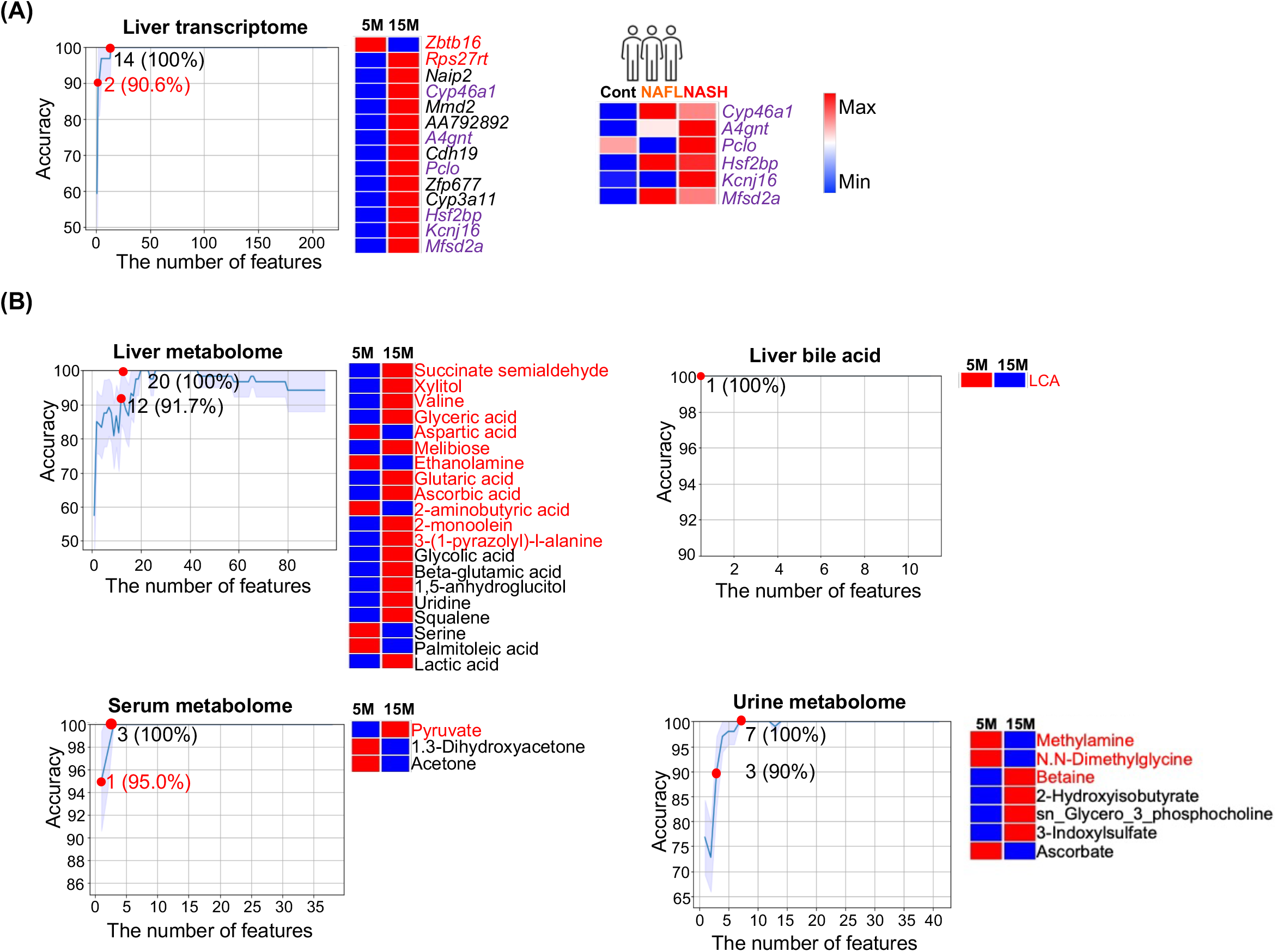
Age classification. The representative line charts show the number of features with the corresponding accuracy in classifying age using (A) liver transcriptome, and (B) metabolomes from the liver, serum, urine as well as hepatic bile acids. (A) Heatmaps show the level of 14 transcripts with 100% accuracy in prediction of age and overlapped transcripts in human NAFLD and NASH cohorts (highlighted in purple). (B) Heatmaps show the differences in the relative abundance of metabolites from liver, serum, and urine as well as hepatic bile acids in 5- and 15-month-old mice (blue and red indicate low and high levels, respectively). The number of features with predictive accuracy > 90% or the number of least features that has the highest accuracy are marked (red dots). The order of features in the heatmap is based on the importance of the feature after feature selection.

The disease relevance of those age-related hepatic transcripts was studied using human datasets. In humans, hepatic *CYP46A1, A4GNT, PCLO, HSF2BP, KCNJ16*, and *MFSD2A* were also found to be elevated in NASH patients compared with healthy controls (Fig. 2A, right panel).

Among the studied molecular signatures including transcripts and metabolites from different sources, hepatic BAs generated the best predicting value to differentiate ages as reduced lithocholic acid (LCA) alone achieved 100% accuracy (Fig. 2B). In addition, twelve liver metabolites (e.g., glyceric acid, melibiose, glutaric acid, etc.), or one serum metabolite (pyruvate), or three urine metabolites (methylamine, N.N-dimethylglycine, and betaine) each had ≥ 90% accuracy in classification of chronological age (Fig. 2).

Integrated pathway analysis was performed for age predictors including transcripts and metabolites (Fig. 3SA). The top regulated pathways are ABC transporters (hepatic aspartic acid, valine, xylitol, uridine, and urine betaine), as well as glycine, serine, and threonine metabolism (hepatic glyceric acid and aspartic acid, urine betaine, and serum pyruvate) (Fig. 3SA).

The disease relevance is elucidated by the metabolite-disease interaction network. In humans, most of the uncovered age-related metabolites were implicated in schizophrenia, Alzheimer’s disease, and lung cancer (Fig. 3SB).

Correlation analysis showed that hepatic LCA was positively associated with serum concentrations of acetone and 1,3-dihydroxyacetone, but negatively correlated with serum pyruvate (Fig. 4S). Instead of using liver samples, our data revealed that serum metabolites (pyruvate, acetone, and 1,3-dihydroxyacetone) are significant in classifying chronological age.

### Predictors for FXR inactivation

FXR whole body KO mice develop NAFL, NASH, and liver tumors spontaneously as they age [27]. WD intake facilitates the progression of disease development [1, 2]. Thus, the inactivation of FXR leads to carcinogenesis within the experimental time frame (i.e., 15 months) [6]. Among the studied groups, 15-month-old WD-fed FXR KO male mice had the most severe hepatic phenotypes, as many of them not only had NASH but also liver tumors [6].

For FXR inactivation, 100% accuracy could be achieved based on the expression pattern of two transcripts (upregulated *Acmsd* and downregulated *Tdg)* or ten hepatic metabolites shown in the heatmap (Fig. 3A, B). Among ten hepatic metabolites, decreased melibiose had 95.8% accuracy to predict FXR inactivation. However, ten hepatic BAs only gave 71.3% accuracy to predict FXR status. Moreover, twelve serum metabolites (succinate, malate, alanine, glutamine, acetone, phenylalanine, methionine, sn-Glycero-3-phosphocholine, urea, glycolate, tyrosine, valine, 2-hydroxyisobutyrate, glucose, 3-hydroxyisobutyrate) predicted FXR expression status with 91.3% accuracy (Fig. 3B). Further, urine creatinine, taurine, and succinate had 95.4% accuracy in predicting FXR status (Fig. 3B). Notably, cecal microbiota at phylum, class, order, family, and genus levels could differentiate FXR KO and WT with > 90% accuracy. Especially, increased *Dorea, Dehalobacterium*, and *Oscillospira* at the genus level yielded 92.7% accuracy to differentiate WT *vs.* FXR KO (Fig. 3C).

**Fig. 3.**
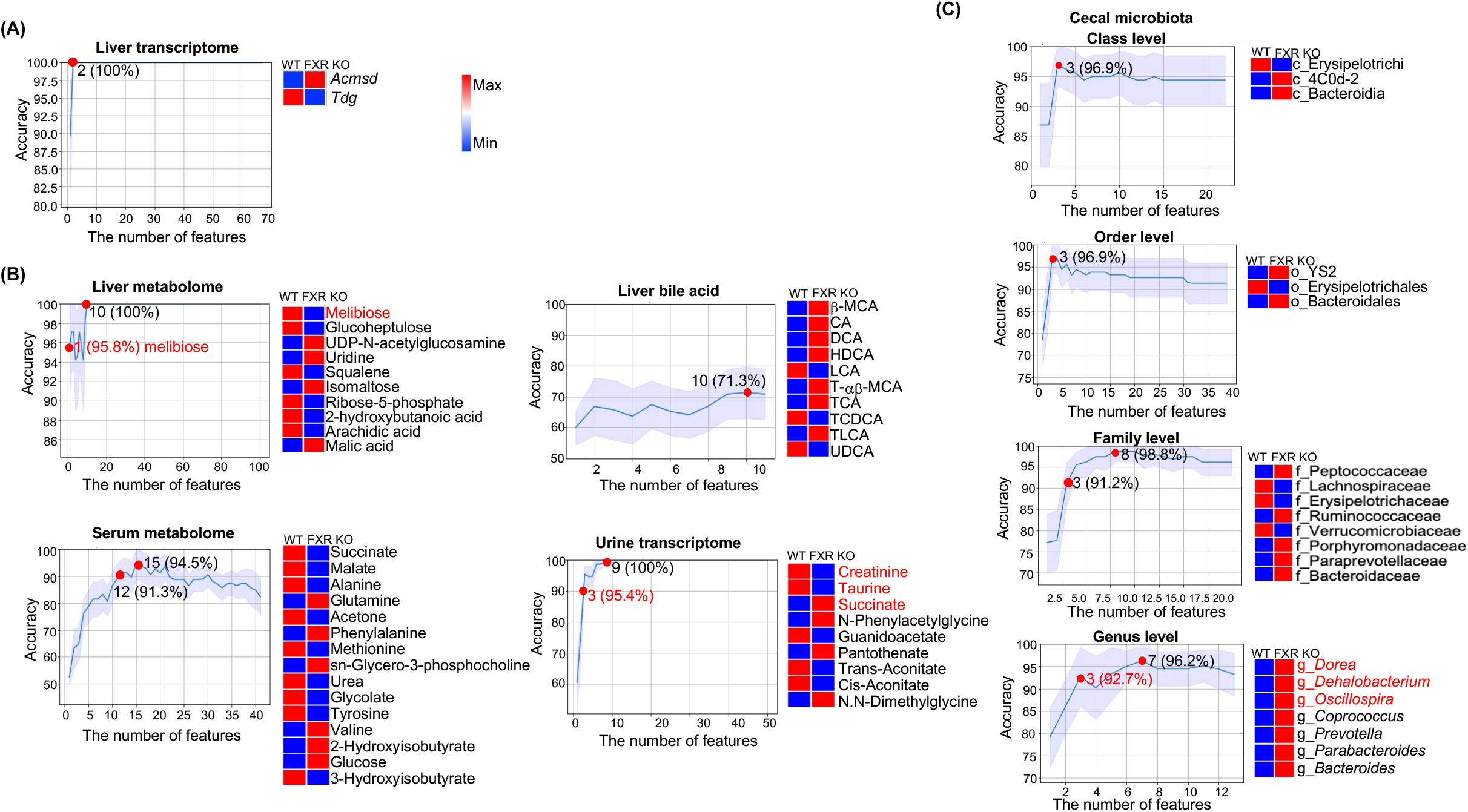
Predictors for FXR expression. Representative line charts show the number of features with indicated accuracy in the prediction of FXR expression status using (A) liver transcriptomes, (B) metabolomes from the liver, serum, and urine as well as hepatic bile acids, and (C) cecal microbiota at different levels. The number of features with predictive accuracy > 90% and/or the number of least features that has the highest accuracy is highlighted (the red dot). The differences in the relative abundance of predictors between FXR KO and WT groups are shown in heatmaps (blue and red indicate low and high levels, respectively). The order of features in the heatmap is based on the importance of the feature after feature selection.

Pathway analysis for metabolites shown in Fig. 3B revealed that serum glutamine, succinate, malate, phenylalanine, methionine, valine, tyrosine, and alanine were involved in the central carbon metabolism in cancer (Fig. 5SA). The metabolite-disease interaction network showed that urine creatinine, which was reduced due to FXR inactivation was associated with neurological disorders (e.g., Canavan disease and schizophrenia), urinary disorders (e.g., Bartter syndrome, type 2, antenatal and Maple syrup urine disease), and metabolic disorders (dimethylglycine dehydrogenase deficiency) (Fig. 5SB). In addition, succinate (succinic acid) was also related to Canavan disease (Fig. 5SB). Urine taurine was associated with Maple syrup urine disease (Fig. 5SB).

Association analysis found that hepatic melibiose was negatively associated with cecal *Dorea, Dehalobacterium*, and *Oscillospira* (Fig. 6SB). Additionally, these three bacteria were also negatively associated with hepatic *Tdg* but positively correlated with hepatic *Acmsd*. It indicates that the increased relative abundance of cecal *Dorea, Dehalobacterium*, and *Oscillospira* can be a marker of FXR inactivity. The roles of FXR status predictors (transcripts and metabolites) are summarized in Table S2-S3.

## Discussion

Our data revealed that the performance of multi-omics in each risk prediction model is different based on the predictive accuracy and the number of features (Table 1). Remarkably, urine metabolite (sucrose), serum metabolites (pyruvate, acetone, and 1,3-dihydroxyacetone), and gut bacteria (*Dorea, Dehalobacterium*, and *Oscillospira)* can classify (>90% accuracy) dietary patterns, ages, and FXR functional status, respectively. The molecular features that act as metabolic liver disease risk predictors are not only biomarkers for the mouse models but also related to human diseases. The information is summarized in Table S2 and S3.

### Diet predictors in metabolic liver disease development

Among the diet predictors, downregulation of *Cyp39a1* (24-hydroxycholesterol 7-alpha-hydroxylase) by WD has been proposed as a novel biomarker for poor overall survival of HCC patients [28]. In contrast, *Gramd1b* (GRAM domain containing 1B), a cholesterol transporter, was upregulated in WD-fed mouse livers suggesting cholesterol overload. Consistent with our findings, the expression of hepatic *Gramd1b* is also increased by a high-cholesterol diet, and silencing hepatic *Gramd1b* in mice suppresses NASH progression [29].

Among the metabolites, reduced hepatic 1,5-anhydroglucitol (an anhydro sugar of D-glucitol) and linoleic acid could predict WD intake with 93.8% accuracy. The 1,5-anhydroglucitol, derived mainly from nearly all foods, is lower in fibrosis stage F3 than in the F0-2 stage in NAFLD patients [30]. The concentration of linoleic acid is also decreased in human HCC tissues compared with normal controls [31]. Linoleic acid is the most abundant ω-6 polyunsaturated fatty acid in human diets, human plasma, and membrane lipids [32].

To develop noninvasive biomarkers of metabolic liver disease risks, we detected urine metabolites and identified that an increase in urine sucrose could be used to predict WD intake. This is not surprising, as the used WD in our animal experiments contains 37% sucrose. It has been shown that there is a significant correlation between the average urinary sucrose excretion and dietary sucrose intake because of the sucrose permeability [33].

### Features that classify ages and metabolic liver diseases

Aging is an inevitable risk factor for most chronic diseases, as it decreases regenerative ability and metabolic processes [34]. *Zbtb16* (zinc finger and BTB domain-containing protein 16), a transcription factor and energy metabolism regulator, is downregulated in aged mice. The *Zbtb16* encoded protein is important in adipogenesis and the control of hepatic gluconeogenesis [35]. In humans, decreased *Zbtb16* variants are associated with elevated total and low-density lipoprotein cholesterol in a sex-dependent manner [36].

Age also affects the profile of BAs, which have pivotal roles in metabolism, immunity, and anti-tumorigenesis. Notably, decreased hepatic LCA could predict older age with 100% accuracy. Consistently, LCA has been identified as an anti-aging compound that extends the lifespan of yeast [37]. LCA acts as an agonist of G-protein-coupled BA receptor TGR5 in increasing free fatty acid availability through lipolysis and induces mitochondrial fission [38]. As the expression of FXR and TGR5 declines with age, dual agonists for FXR and TGR5 have been shown to delay age-related kidney deterioration in mouse models [39]. In humans, isoforms of LCA (iso-, 3-oxo-, allo-, 3-oxoallo- and isoallolithocholic acid)-producing bacteria were enriched in centenarians [40]. In rats, dietary conjugated LCA, a mixture of positional and geometric isomers of linoleic acid, alleviates NAFLD [41]. Taken together, LCA may be a target for aging related NAFLD treatment.

Our data revealed that serum pyruvate as well as acetone (a ketone body) and 1,3-dihydroxyacetone (DHA) correlated with hepatic LCA. The potential of reduced serum pyruvate together with increased serum acetone and DHA being a metabolically active young liver, warrants further validation in humans. Serum pyruvate is derived from alanine and a-ketoglutarate converted by the alanine aminotransferase (ALT) and elevated ALT is a diagnostic marker for liver injury. The concentration of serum pyruvate was also elevated by high-fat diet intake [42]. 1,3-Dihydroxyacetone is a 3-carbon reducing sugar produced from glycerol. Acetone is the simplest ketone body and is synthesized from fatty acid oxidation in the livers. Thus, reduced serum acetone likely indicates reduced fatty acid oxidation. Moreover, elevated breath acetone is a biomarker of type 2 diabetes mellitus in the breath analysis [43]. Whether reduced serum acetone can be a biomarker for reduced fatty acid oxidation associated with aging liver also warrants further investigation.

### FXR inactivation predictors and metabolic liver diseases

Hepatic transcripts *Acmsd* (aminocarboxymuconate semialdehyde decarboxylase) and *Tdg* (G/T mismatch-specific thymine DNA glycosylase) could differentiate FXR KO from WT. Upregulated *Acmsd* and downregulated *Tdg* in the livers were signatures of FXR inactivation. ACMSD controls cellular NAD+ levels in the liver [44]. Inhibition of *Acmsd* attenuates hepatic steatosis and reduces liver injury in diet-induced NAFLD mouse models [45]. TDG (thymine DNA glycosylase) is an enzyme that plays a key role in active DNA demethylation. It is essential for maintaining glucose and BA homeostasis, as depletion of *Tdg* causes dysregulation of FXR signaling and leads to HCC development in mice [46].

It is interesting to note that the increased abundance of *Dorea, Dehalobacterium*, and *Oscillospira* in cecal content has greater than 90% accuracy in FXR KO prediction. In humans, the abundance of *Dorea* is also increased in NAFLD patients compared with healthy controls [47]. *Dehalobacterium* is known to have a negative association with the body mass index [48]. *Oscillospira* is increased in high-fat diet-fed mice compared with normal controls [49].

Urine metabolites also predicted FXR functional status. As a signature of FXR KO, urine creatinine and taurine decreased while succinate increased. Urine creatinine reflects muscle mass, and low urine creatinine is associated with cardiovascular disease risk [50]. Taurine is beneficial in alleviating fatty liver disease by promoting energy expenditure and preventing oxidative damage and inflammation [51]. Succinate is an inflammation-induced immunoregulatory metabolite in the macrophages [52], it is also elevated in the inflammation [53]. Thus, the metabolic features that predict FXR inactivation are involved in metabolism and immune responses.

Collectively, the study has identified features from different sources that have different predicting power to differentiate risks for metabolic disease development. Urine or gut microbiota biomarkers can be valuable for noninvasive diagnosis of liver function status. As WD intake, aging, and FXR inactivation are also implicated in other diseases including dermatitis and dementia, the uncovered risk predictors have multiple disease implications [8–10].

## Abbreviations

*Acmsd*: aminocarboxymuconate semialdehyde decarboxylase
*Adam11*: ADAM metallopeptidase domain 11
*A4gnt*: alpha-1,4-N-acetylglucosaminyltransferase
*CD*: control diet
*Cdh19*: cadherin 19
*Cyp39a1*: cytochrome P450 family 39 subfamily a member 1
*Cyp3a11*: cytochrome P450 family 3 subfamily a member
*Cyp46a1*: cytochrome P450 family 46 subfamily a member 1
*Csad*: cysteine sulfinic acid decarboxylase
*DEGs*: differentially expressed genes
*FXR*: farnesoid X receptor
*GEO*: Gene Expression Omnibus
*Gramd1b*: GRAM domain containing 1B
*Hamp2*: hepcidin antimicrobial peptide 2
*HCC*: hepatocellular carcinoma
*Hsf2bp*: heat shock transcription factor 2 binding protein
*Kcnj16*: potassium inwardly rectifying channel subfamily J member 16
*Loxl4*: lysyl oxidase like 4
*ML*: machine learning
*Mfsd2a*: major facilitator superfamily domain containing 2A
*Mmd2*: monocyte to macrophage differentiation associated 2
*NAFLD*: nonalcoholic fatty liver disease
*NAFL*: nonalcoholic fatty liver
*NASH*: non-alcoholic steatohepatitis
*Naip2*: NLR family apoptosis inhibitory protein
*Pclo*: piccolo presynaptic cytomatrix protein
*Pde5a*: phosphodiesterase 5A
*Rec8*: REC8 meiotic recombination protein
*Rps27rt*: Ribosomal Protein S27
*SVM*: support vector machine
*Slc39a4*: solute carrier family 39 member 4
*Tdg*: thymine DNA gycosylase
*WD*: Western diet
*Zbtb16*: zinc finger and BTB domain containing 16
*Zfp677*: zinc finger protein

## Acknowledgments

We thank Shuai Zhang and Cecylia Olivo for their contributions to preparing and editing the manuscript.

## Authors’ contributions

YJW and GYY conceived and designed the study. GYY and RL analyzed data; XL and RS participated in machine learning application and manuscript preparation; GYY and YJW wrote the manuscript; all authors commented and approved the final manuscript.

## Disclaimers

The findings and conclusions in this report are those of the authors and do not necessarily represent the views or opinions of the California Department of Public Health or the California Health and Human Services Agency. The data were generated using a specific number of a certain strain of mice, which is standard for basic research. Whether the findings apply to all animal species requires validation.

## Data and code availability

Hepatic bulk RNA sequencing data are available on the GEO database (https://www.ncbi.nlm.nih.gov/geo/; GSE216375). Bioinformatics and statistical results for hepatic transcriptome used for feature selection were available in a previous study [16]. All python scripts used, and additional information related to this paper can be requested from the authors.

## Supplementary Information

### Additional file 1

**Table S1**. The sample information of multi-omics data that used for training and validation.

### Additional file 2

**Fig. 1S. Functional analysis of diet predictors.** (A) Integrated pathway analysis showing pathways for WD-predictors (transcripts and metabolites). The corresponding features for the important pathways are indicated. (B) The network shows that metabolomic predictors of WD intake are associated with human diseases.

**Fig. 2S. Spearman’s correlation for WD-predictors from the liver, serum, and urine.** Spearman’s correlation, \**p* < 0.05, \*\**p* < 0.01.

**Fig. 3S. Functional analysis of age-predictors.** (A) Integrated pathway analysis for age-predictors (metabolites). (B) Features that can classify ages in association with human diseases.

**Fig. 4S. Interaction between features that can be used for chronological age prediction.** Spearman’s correlation, *\*p* < 0.05, \*\**p* < 0.01.

**Fig. 5S. Functional analysis of FXR expression predictors.** (A) The pathways for metabolites serve as FXR expression predictors. (B) The network shows the interaction between metabolites and diseases for FXR expression predictors.

**Fig. 6S. Interactions of FXR expression predictors.** Spearman’s correlation between cecal microbiota at the genus level, hepatic transcripts, and metabolites from the liver, serum, and urine. \**p* < 0.05, \*\**p* < 0.01.

### Additional file 3

**Table S2**. Summary table of transcriptomic predictors of diet, age, and FXR expression

### Additional file 4

**Table S3**. Summary table of metabolomic predictors of diet, age, and FXR expression

