## Additional file 1 for "Machine Learning to Identify Molecular Markers for Metabolic Disease Development Using Mouse Models"

**Table S1. Samples for multi-omics data from the mouse model**

| Omics | Number of mice | Age (month) | Diet | Genotype |
|---|---|---|---|---|
| **Hepatic transcripts** | 48 | 5, 10, 15 | CD, WD | WT, FXR KO |
| **Metabolites** | | | | |
| Bile acids (liver) | 186 | 5, 10, 15 | CD, WD | WT, FXR KO |
| Liver | 72 | 5, 10, 15 | CD, WD | WT, FXR KO |
| Serum | 122 | 5, 10, 15 | CD, WD | WT, FXR KO |
| Urine | 157 | 5, 10, 15 | CD, WD | WT, FXR KO |
| **Microbiota** | 163 | 5, 10, 15 | CD, WD | WT, FXR KO |
