## Additional file 2 for "Machine Learning to Identify Molecular Markers for Metabolic Disease Development Using Mouse Models"

**Fig. 1S Diet prediction**

**(A)**

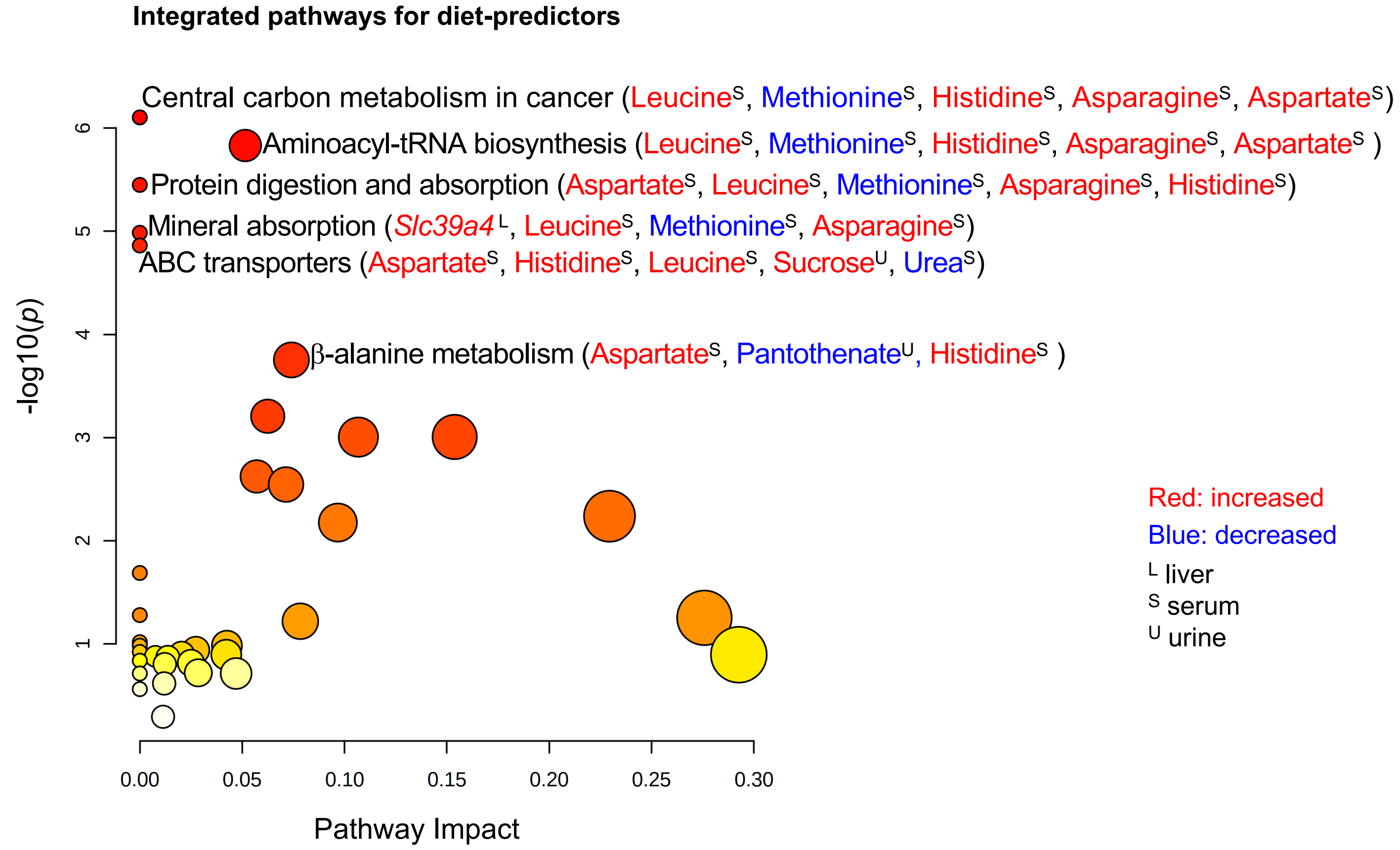

**Fig. 1S Diet prediction**

**(B)**

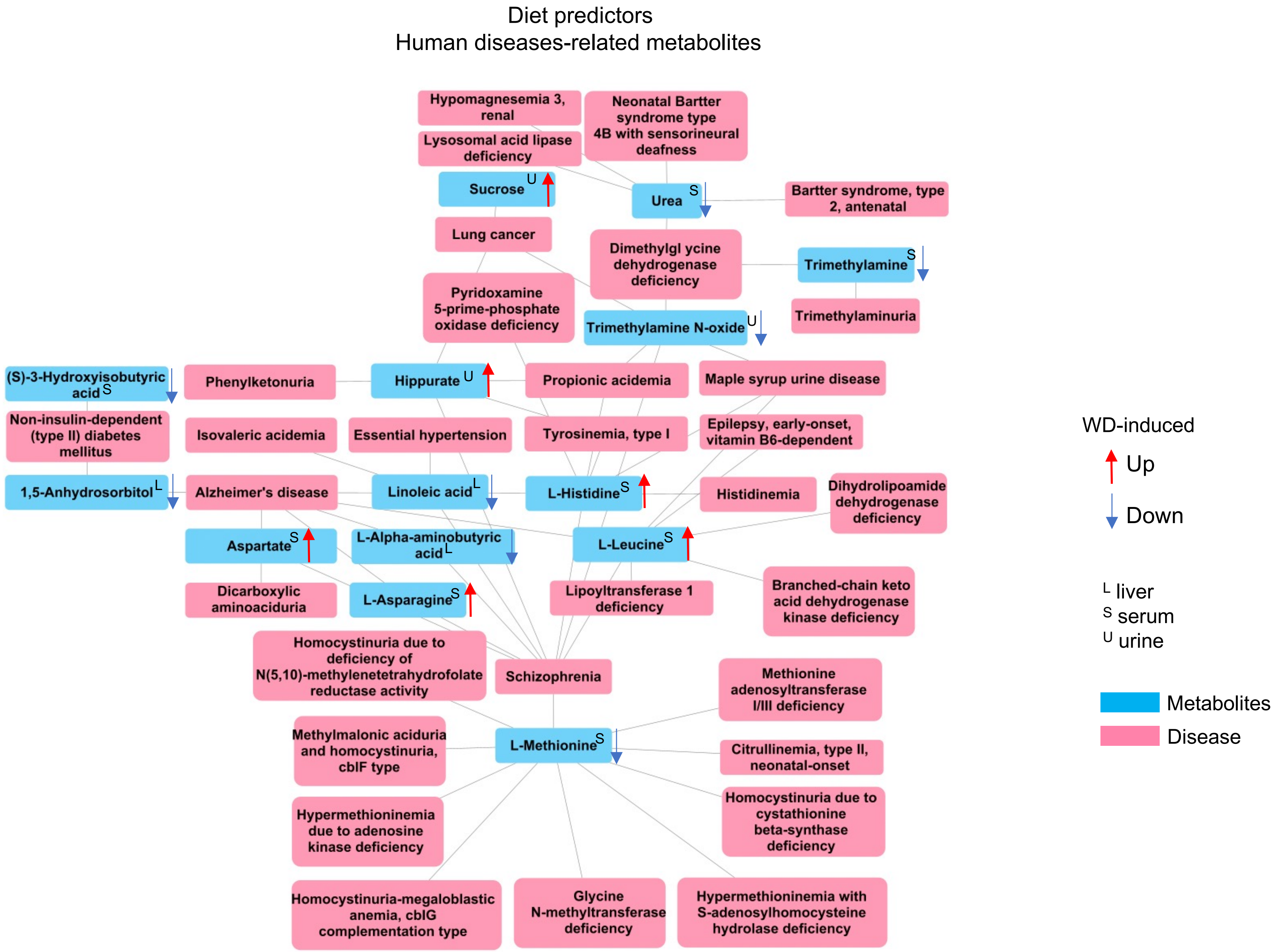

**Fig. 2S Spearman's correlation between diet-predictors**

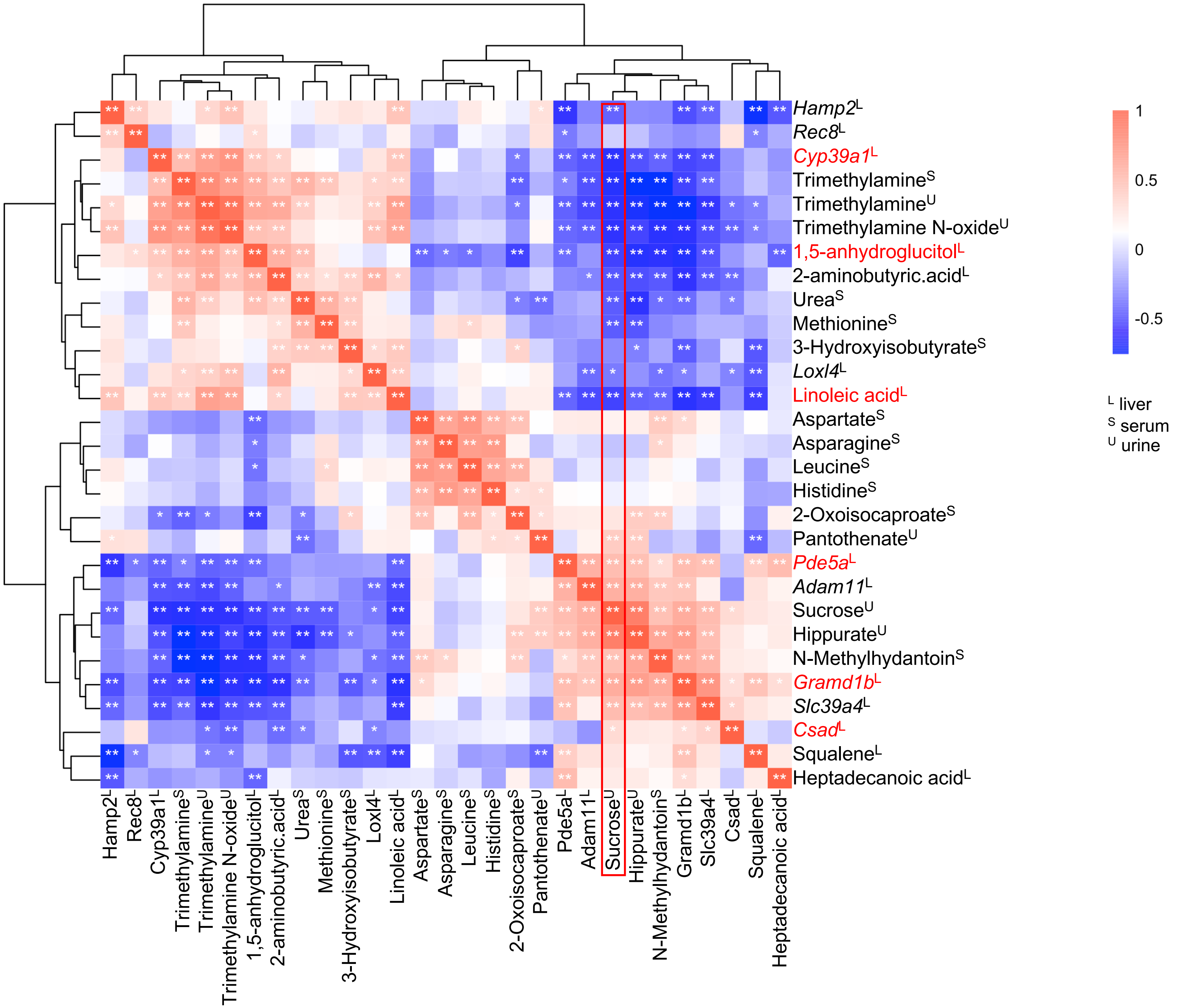

**Fig. 3S Chronological age prediction**

**(A)**

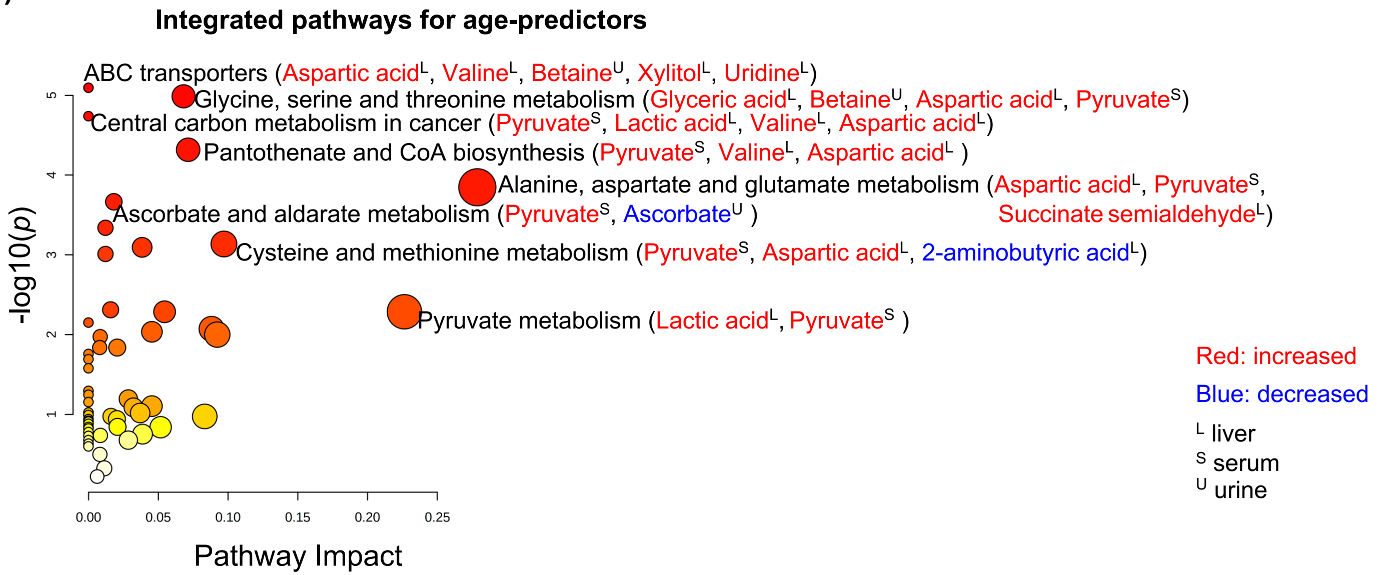

**Fig. 3S Chronological age prediction**

**(B)**

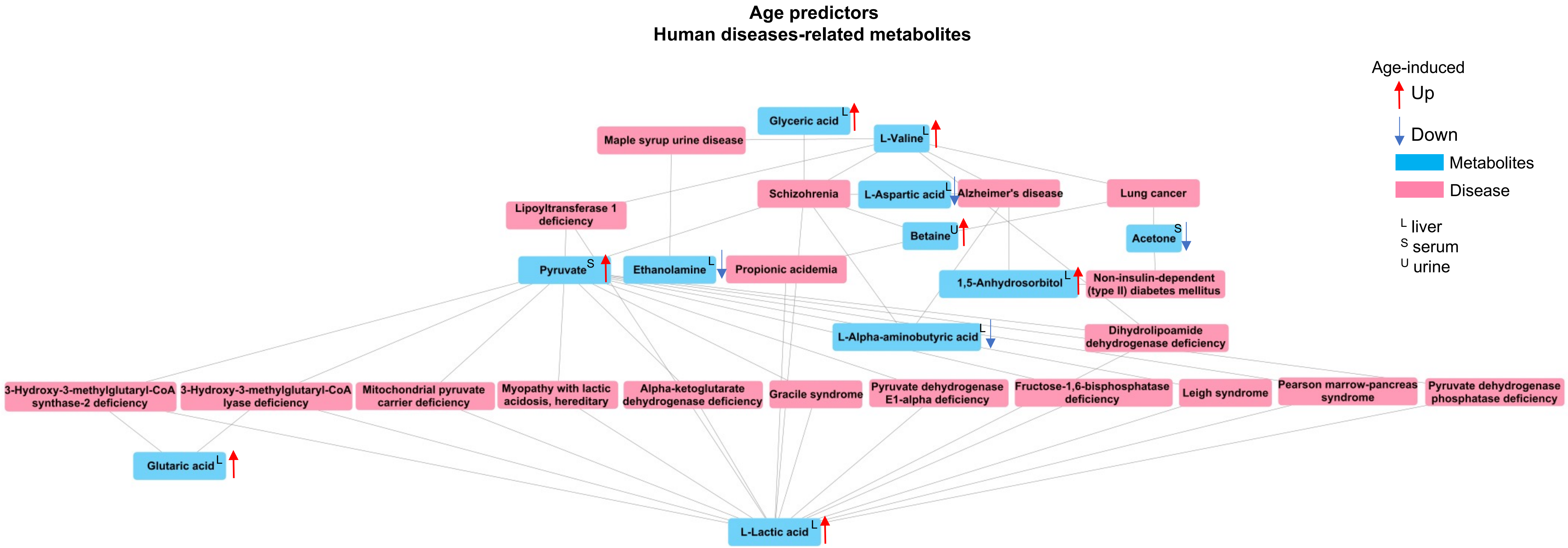

**Fig. 4S Spearman's correlation between age-predictors**

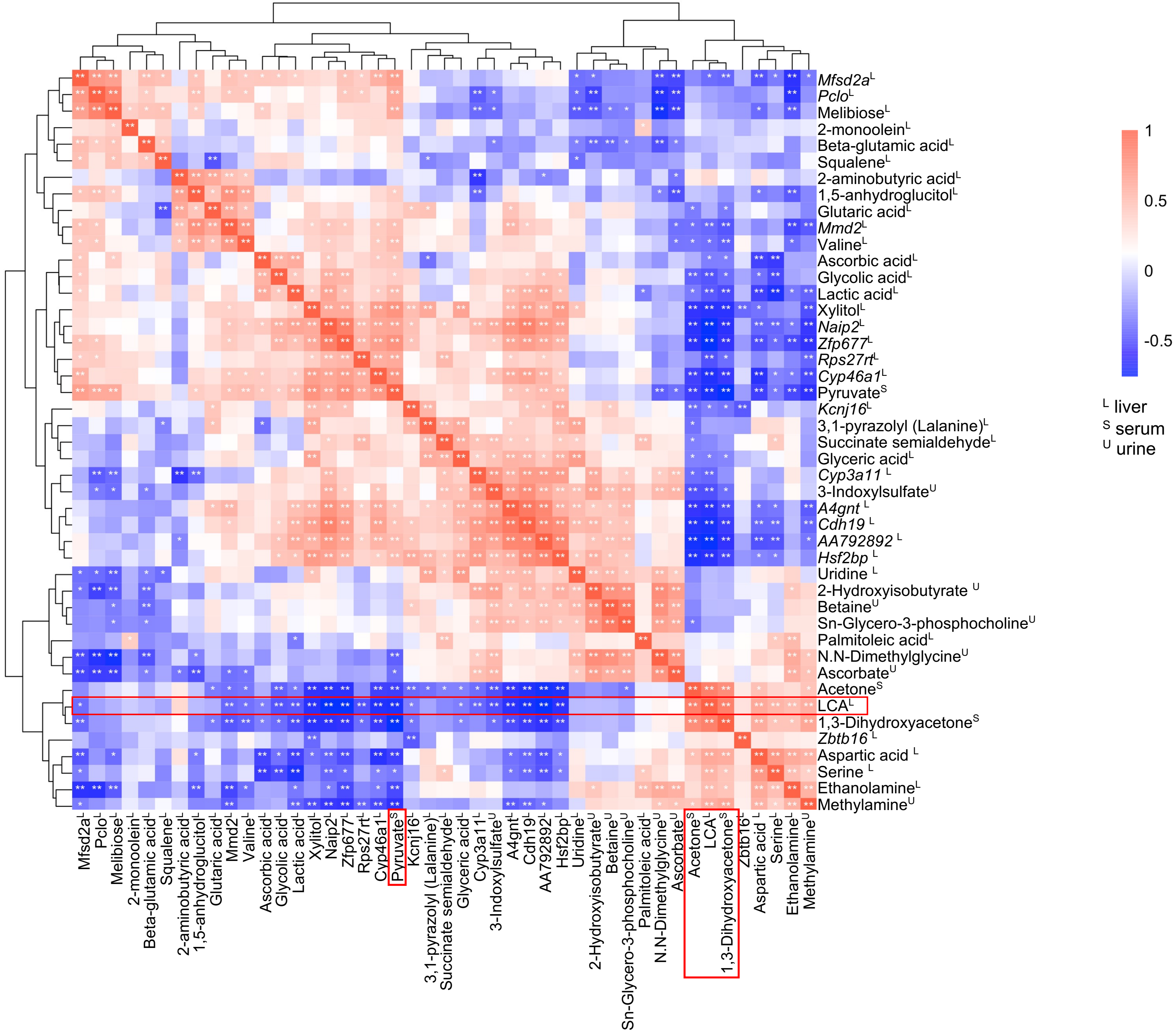

**Fig. 5S FXR deactivation prediction**

**(A)**

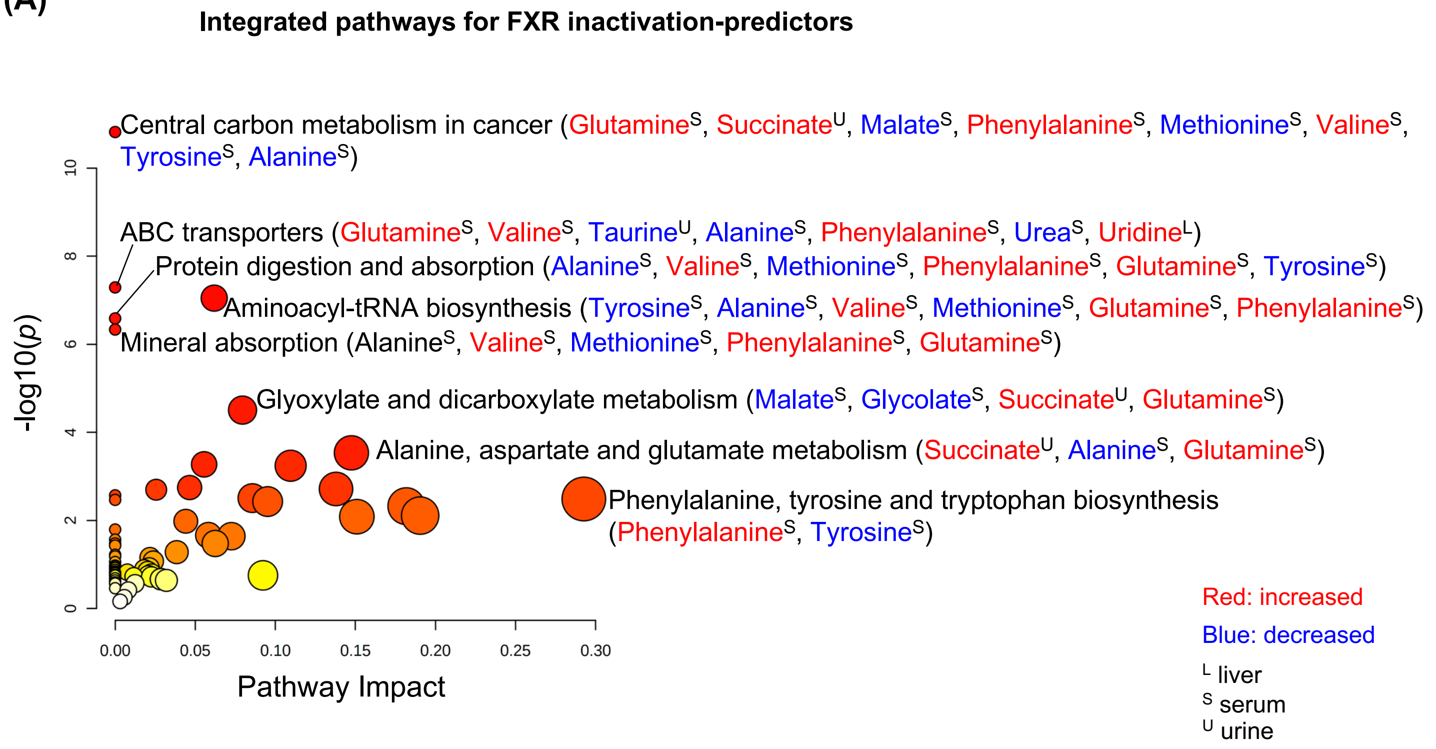

**Fig. 5S FXR deactivation prediction**

**(B)**

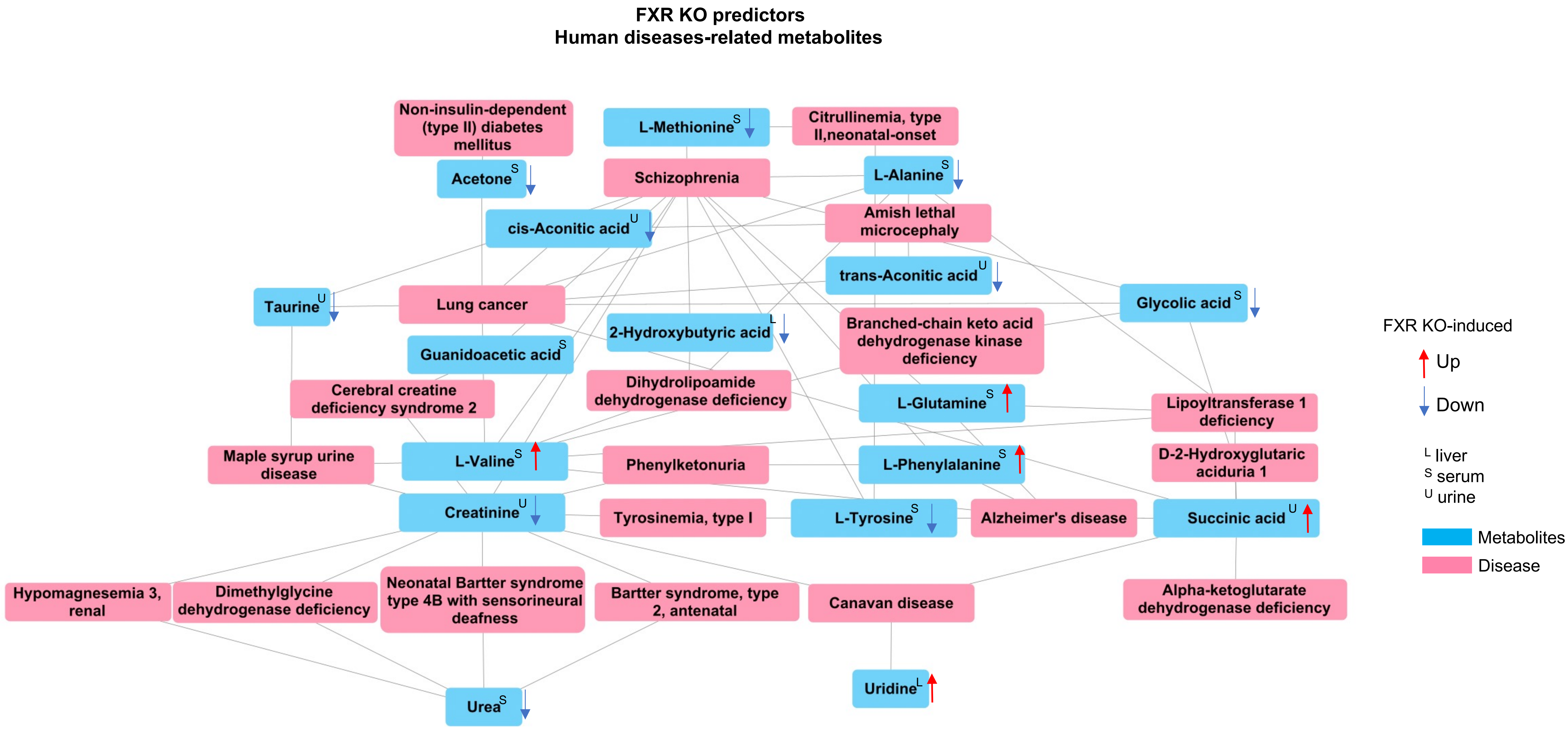

Fig. 6S Spearman's correlation between FXR KO-predictors

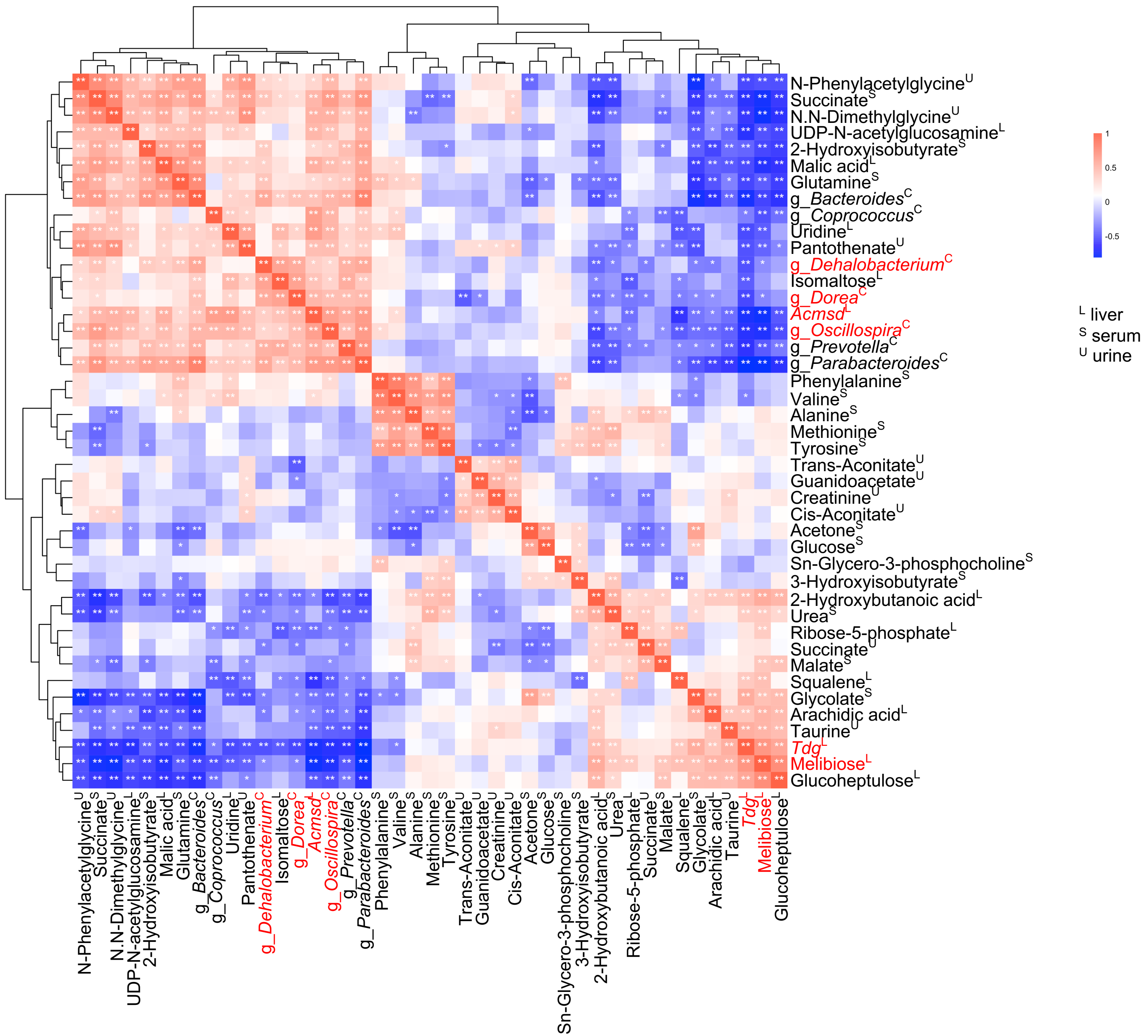
