## Additional file 3 for "Machine Learning to Identify Molecular Markers for Metabolic Disease Development Using Mouse Models"

**Table S2. Hepatic transcripts that classify diet, age, and FXR activity**

| Transcript name | Protein name | Functions | Changes | Diseases implications |
|---|---|---|---|---|
| *Cyp39a1* | 24-hydroxycholesterol 7-alpha-hydroxylase | Steroid metabolism; cholesterol degradation; lipid metabolism; bile acid biosynthesis | Western diet (Down) | CYP39A1 is an HCC suppressor in humans [1].<br>CYP39A1 is an HCC biomarker [2]. |
| *Pde5a* | cGMP-specific 3',5'-cyclic phosphodiesterase | Purine metabolism, plays a role in signal transduction by regulating the intracellular concentration of cyclic nucleotides | Western diet (Up) | - |
| *Csad* | Cysteine sulfinic acid decarboxylase | Organosulfur biosynthesis; taurine biosynthesis | Western diet (Up) | CSAD is protective in NAFLD [3]. |
| *Gramd1b* | Protein Aster-B | Cholesterol transporter that mediates non-vesicular transport of cholesterol from the plasma membrane to the endoplasmic reticulum | Western diet (Up) | - |
| *Slc39a4* | Zinc transporter ZIP4 | ZIP4 (SLC39A4) may play a role in the acquisition of zinc by hepatocellular carcinomas, leading to repressed apoptosis, enhanced growth rate and enhanced invasive behavior [4]. | Western diet (Up) | SLC39A4 expression is activated in HCC [4].<br>SLC39A4 is involved in the pathogenesis of acrodermatitis enteropathica, zinc-deficiency type [5-7]. |
| *Hamp2* | Hepcidin-2 | Antimicrobial | Western diet (Down) | - |

| | | | | |
|---|---|---|---|---|
| *Loxl4* | Lysyl oxidase homolog 4 | Crosslinking of collagen fibrils; Elastic fibre formation | Western diet (Down) | LOXL4 expression is increased during the liver carcinogenesis [8]. |
| *Rec8* | Meiotic recombination protein REC8 homolog | Meiotic synapsis | Western diet (Down) | REC8 promotes tumor migration, invasion, and angiogenesis in human HCC [9]. |
| *Adam11* | Disintegrin and metalloproteinase domain-containing protein 11 | LGI (leucine-rich glioma inactivated)-ADAM interactions | Western diet (Up) | - |
| *Zbtb16* | Zinc finger and BTB domain-containing protein 16 | Energy metabolism regulator | Aging (Down) | ZBTB16 associate with increased obesity-related parameters and elevated total and low-density lipoprotein cholesterol [10]. |
| *Rps27rt* | Ribosomal Protein S27 | RNA binding and structural constituent of ribosome | Aging (Up) | - |
| *Naip2* | Baculoviral IAP repeat-containing protein 1b | Inflammatory response; Innate immunity; Apoptosis | Aging (Up) | - |
| *Cyp46a1* | Cholesterol 24-hydroxylase | Steroid metabolism; cholesterol degradation; Lipid metabolism; C21-steroid hormone metabolism | Aging (Up) | *Cyp46* gene levels were higher in the hippocampus of elderly humans and in patients with certain pathological, neurodegenerative conditions [11]. |
| *Mmd2* | Monocyte to macrophage differentiation factor 2 | Positive regulation of Ras protein signal transduction | Aging (Up) | - |
| *AA792892* | Expressed sequence AA792892 | | Aging (Up) | - |

| | | | | |
|---|---|---|---|---|
| *A4gnt* | Alpha-1,4-N-acetylglucosaminyltransferase | O-linked glycosylation of mucins | Aging (Up) | - |
| *Cdh19* | Cadherin-19 | Wnt signaling pathway; Cadherin signaling pathway | Aging (Up) | - |
| *Pclo* | Protein piccolo | Component of the presynaptic cytoskeletal matrix, involved in regulation of presynaptic proteins and synaptic vesicles | Aging (Up) | Loss of PCLO causes pontocerebellar hypoplasia 3 in humans [12];<br>*Pclo* is highly mutated as cancer driver gene in hepatitis B virus-related HCC patients [13]. |
| *Zfp677* | Uncharacterized protein | - | Aging (Up) | - |
| *Cyp3a11* | Cytochrome P450 3A11 | Xenobiotics; Aflatoxin activation and detoxification; Biosynthesis of maresin-like SPMs | Aging (Up) | The mRNA expression of hepatic *Cyp3a11 is increased* high-fat and high-sucrose induced NAFLD mouse model [14]. |
| *Hsf2bp* | Heat shock factor 2-binding protein | Double-strand break repair involved in meiotic recombination | Aging (Up) | - |
| *Kcnj16* | Inward rectifier potassium channel 16 | Potassium transport channels; Activation of G protein gated Potassium channels; Inhibition of voltage gated Ca2+ channels via Gbeta/gamma subunits | Aging (Up) | *Kcnj16* level is lower in HCC tumor tissues compared to adjacent tissues from HCC patients [15]. *Kcnj16* mutation cause a novel tubulopathy with hypokalemia, salt wasting, disturbed acid-base homeostasis, and sensorineural deafness [16]. |

| | | | | |
|---|---|---|---|---|
| *Mfsd2a* | Sodium-dependent lysophosphatidylcholine symporter 1 | Glycerophospholipid biosynthesis | Aging (Up) | The MFSD2A level in HCC patients is lower than in healthy controls [17]. |
| *Acmsd* | 2-amino-3-carboxymuconate-6-semialdehyde decarboxylase | Secondary metabolite metabolism; quinolate metabolism | FXR KO (Up) | Inhibiting ACMSD protects liver injury in mouse models [18]. |
| *Tdg* | G/T mismatch-specific thymine DNA glycosylase | DNA demethylation | FXR KO (Down) | Conditional knockout of Tdg causes HCC in a mouse model [19]. |

HCC, hepatocellular carcinoma; NAFLD, nonalcoholic liver disease; -, unknown
