## Additional file 4 for "Machine Learning to Identify Molecular Markers for Metabolic Disease Development Using Mouse Models"

**Table S3. Metabolites that classify diet, age, and FXR activity**

| Metabolites (Specimens) | Source | Functions or Roles | Changes | Disease implications |
|---|---|---|---|---|
| 1,5-anhydroglucitol (Liver) | Mainly from food, well absorbed in the intestine, and is distributed to all organs and tissues. | A glycemic marker | Western diet (Down)<br>Aging (Up) | Serum 1,5-anhydroglucitol concentrations are lower in cirrhotic and chronic liver disease patients than those in healthy people [1, 2].<br>Low serum 1,5-anhydroglucitol concentration is a predictor of short-term mortality in Hepatitis B virus-related acute-on-chronic liver failure patients [3]. |
| Linoleic acid (Liver) | Hydrogenated vegetable oils | An omega-6 trans fatty acid; $\alpha$-linolenic acid and linoleic acid metabolism | Western diet (Down) | Dietary conjugated linoleic acid protects the liver from NAFLD in a rat model [4]. |
| Squalene (Liver) | Human sebum (5%), fish liver oils, yeast lipids, and many vegetable oils (e.g., palm oil, cottonseed oil, rapeseed oil) | A biochemical precursor to the whole family of steroids; An intermediate for cholesterol synthesis; A precursor to phytosterol synthesis; A bactericide | Western diet (Up)<br>Aging (Up)<br>FXR KO (Down) | Squalene decreases hepatic cholesterol and triglycerides [5]. |
| Heptadecanoic acid (Liver) | Exogenous origin, such as dairy fats (milk and meat) | A saturated fatty acid | Western diet (Up) | Biomarkers for dietary food (fat) intake assessment; Biomarkers for coronary heart disease risk and type II diabetes mellitus risk [6]. |
| 2-aminobutyric acid (Liver) | Biosynthesized by transamination of oxobutyrate; | A non-proteinogenic $\alpha$-amino acid | Western diet (Down) | - |

| | | | | |
|---|---|---|---|---|
| | A metabolite in isoleucine biosynthesis; A non-essential amino acid that is primarily derived from the catabolism of methionine, threonine, and serine. | | | |
| Trimethylamine (Serum) | A product of decomposition of plants and animals; A bacteria metabolite | A colorless, hygroscopic, and flammable simple amine with a typical fishy odor in low concentrations; Ammonia like odor in higher concentrations. | Western diet (Down) | NAFLD patients have increased trimethylamine in the intestine and liver [7]. |
| 3-Hydroxyisobutyrate (Serum) | An intermediate in L-valine metabolism | Valine, leucine, and isoleucine degradation | Western diet (Down)<br>FXR KO (Down) | Plasma 3-hydroxyisobutyrate is a marker of hepatic mitochondrial fatty acid oxidation in male Wistar rats [8]. |
| Aspartate (Serum) | Found in all organisms ranging from bacteria to plants to animals; Nonessential amino acid derived from glutamic acid by enzymes using vitamin B6. | Proteinogenic $\alpha$-amino acid; Arginine and proline metabolism; Aspartate metabolism; Urea cycle | Western diet (Up) | Administration of L-aspartate in vitro or in mice efficiently ameliorates metabolic dysfunction-associated fatty liver disease [9]. |
| Leucine (Serum) | Essential amino acid; Human dietary sources are foods that contain protein, such as meats, dairy products, soy products, beans and legumes. | A branched chain amino acid; Proteinogenic $\alpha$-amino acid;<br>Valine, leucine, and isoleucine degradation | Western diet (Up) | L-leucine supplementation is protective in patients with liver cirrhosis [10]. |

| | | | | |
|---|---|---|---|---|
| Histidine (Serum) | Essential amino acid; Exogenous food, such as fruits (berries) | $\alpha$-amino acid; Histidine metabolism; Nitrogen metabolism; Beta-alanine metabolism; Anti-oxidant, anti-inflammatory and anti-secretory properties | Western diet (Up) | Histidine supplementation ameliorates metabolic syndrome [11]. |
| Urea (Serum) | Exogenous food, such as berries | Urea cycle; Arginine and proline metabolism | Western diet (Down)<br>FXR KO (Down) | Urea cycle disorders [12]. |
| 2-Oxoisocaproate (Serum) | Endogenously produced metabolite | Valine, leucine, and isoleucine degradation; A neurotoxin and a metabotoxin. | Western diet (Up) | Plasma 2-oxoisocaproate (ketoleucine) concentrations are increased in patients with metabolic stress [13]. |
| N-Methylhydantoin (Serum) | Exogenous, such as animal foods. | A bacterial metabolite; A imidazolidine-2,4-dione that is the N-methyl-derivative of hydantoin. | Western diet (Up) | - |
| Methionine (Serum) | Essential amino acid; Exogenous food such as berries; An intermediate in transmethylation reactions | Proteinogenic $\alpha$-amino acid; Glycine, serine, and threonine metabolism; Methionine metabolism | Western diet (Down)<br>FXR KO (Down) | NAFLD patients have low liver methionine concentrations [14]. |
| Asparagine (Serum) | Non-essential amino acid, the precursor to asparagine is oxaloacetate; Exogenous food such as berries | Proteinogenic $\alpha$-amino acid; Aspartate metabolism; Nitrogen | Western diet (Up) | Upregulating asparagine synthetase is beneficial for alleviating liver injury [15]. |

| | | | | |
|---|---|---|---|---|
| | | metabolism; Ammonia recycling | | |
| Sucrose (Urine) | From sugarcane (Saccharum officinarum), sugar beet (Beta vulgaris), and other plants | Sucrose is a sweetener and used in food products as a preservative, antioxidant, moisture control agent, stabilizer, and thickening agent; Galactose metabolism | Western diet (Up) | Excessive sucrose intake leads to NAFLD [16]. |
| Trimethylamine (Urine) | Decomposition of plants and animals | A uremic toxin; A marker for urinary tract infection brought on by *E. coli*. | Western diet (Down) | Trimethylamine in the intestine and liver is increased in NAFLD patients [7]. |
| Trimethylamine N-oxide (Urine) | Biosynthesized endogenously from trimethylamine, which is derived from choline. | A uremic toxin; TMAO alters cholesterol metabolism in the intestines, in the liver and in arterial wall. | Western diet (Down) | Trimethylamine-N-oxide promotes brain aging and cognitive impairment in mice [17]. TMAO supplementation aggravates liver steatosis [18]. |
| Hippurate (Urine) | Produced from the metabolism of benzoate, which is mainly stored in the liver mitochondria. | Reflect hepatic function | Western diet (Up) | Elevated blood hippurate associates with improved hepatic steatosis and good metabolic health [19]. |
| Pantothenate Vitamin B5 (Urine) | From everywhere and small quantities of pantothenic acid are found in nearly every food, with high amounts in whole | A water-soluble vitamin required to sustain life; Beta-alanine metabolism; Pantothenate and CoA biosynthesis | Western diet (Down) FXR KO (Up) | - |

| | | | | |
|---|---|---|---|---|
| | grain cereals, legumes, eggs, meat, and royal jelly. | | | |
| Succinate semialdehyde (Liver) | An intermediate in the catabolism of gamma-aminobutyrate or gamma-aminobutyric acid | High levels of succinate semialdehyde function as a neurotoxin and a metabotoxin. | Aging (Up) | - |
| Xylitol (Liver) | Xylitol exists in all living species, ranging from bacteria to plants to humans. | Organism's growth, development, or reproduction | Aging (Up) | - |
| Valine (Liver) | Essential amino acid; Human dietary sources are foods that contain protein, such as meats, dairy products, soy products, beans and legumes. | Proteinogenic $\alpha$-amino acid; Branched chain amino acid; Valine, leucine, and isoleucine degradation; Propanoate metabolism; Transcription/Translation | Aging (Up) | Excessive valine causes NAFLD [20]. |
| Glyceric acid (Liver) | Obtained from oxidation of glycerol. | Glycine, serine, and threonine metabolism; Glycerolipid metabolism | Aging (Up) | Oral D-glyceric acid activates mitochondrial metabolism and reduces inflammation among 50-60-year-old healthy volunteers [21]. |
| Aspartic acid (Liver) | Aspartic acid is found in all organisms ranging from bacteria to plants to animals. | Proteinogenic $\alpha$-amino acid; Transcription/translation; Arginine and proline metabolism; | Aging (Down) | - |

| | | | | |
|---|---|---|---|---|
| | | Aspartate metabolism | | |
| Ethanolamine (Liver) | Ethanolamine exists in all living species, ranging from bacteria to plants to humans. | An initial precursor for the biosynthesis of two primary phospholipid classes, phosphatidylcholine (PC) and phosphatidylethanolamine (PE). | Aging (Down) | Ethanolamine protects against hyperlipidemia in aged mice [22]. |
| Glutaric acid (Liver) | Glutaric acid is naturally produced in the body during the metabolism of some amino acids, including lysine and tryptophan. | Glutaric acid may cause irritation to the skin and eyes. Glutaric acid acts as an acidogen and a metabotoxin when present in sufficiently high levels. | Aging (Up) | - |
| Ascorbic acid (Liver) | Ascorbic acid is found naturally in citrus fruits and many vegetables and is an essential nutrient in human diets. | Necessary to maintain connective tissue and bone.<br>Tyrosine metabolism | Aging (Up) | Ascorbic acid inhibits obesity and nonalcoholic fatty liver disease [23]. |
| 2-aminobutyric acid (Liver) | A non-essential amino acid; Biosynthesized by transamination of oxobutyrate, a metabolite in isoleucine biosynthesis. | Non-proteogenic amino acid | Aging (Down) | - |

| | | | | |
|---|---|---|---|---|
| 2-monoolein (Liver) | A major end product of the intestinal digestion of dietary fats in animals via the enzyme pancreatic lipase. | Act as emulsifiers, helping to mix ingredients such as oil and water that would not otherwise blend well | Aging (Up) | - |
| 3-(1-pyrazolyl)-l-alanine (Liver) | L-alanine derivative | A non-proteinogenic L-$\alpha$-amino acid | Aging (Up) | - |
| Glycolic acid (Liver, serum) | A metabolite in bacteria such as Acetobacter, *Escherichia* | A nephrotoxin if consumed orally; A known inhibitor of tyrosinase; Renal toxicity | Aging (Up in the liver)<br>FXR KO (Down in serum) | - |
| Beta-glutamic acid (Liver) | A metabolite found in the aging mouse brain; A natural product found in Chondria armata | A marine metabolite and an algal metabolite. | Aging (Up) | - |
| Uridine (Liver) | Uridine can be synthesized from uracil. Uridine is found in many foods (anything containing RNA). | A nucleoside consisting of uracil and D-ribose and a component of RNA; Pyrimidine metabolism | Aging (Up)<br>FXR KO (Up) | Uridine alleviates carbon tetrachloride-induced liver fibrosis [24]. |
| Serine (Liver) | Serine is found in all organisms ranging from bacteria to plants to animals. | Proteinogenic amino acid; Nitrogen metabolism; Starch and sucrose metabolism;<br>Glycine, serine, and threonine metabolism | Aging (Down) | Reduced hepatic serine contributes to the development of fatty liver disease [25]. |

| | | | | |
|---|---|---|---|---|
| Palmitoleic acid (Liver) | A common constituent of the glycerides of human adipose tissue; From gut bacteria, such as Akkermansia muciniphila; Macadamia oil (Macadamia integrifolia) and sea buckthorn oil (Hippophae rhamnoides) are botanical sources of palmitoleic acid, containing 22 and 40% respectively. | A monounsaturated fatty acid; Reduce hepatic gluconeogenesis | Aging (Down) | Palmitoleic acid reduces high fat diet-induced liver inflammation [26]. Palmitoleic acid improves metabolic functions in fatty liver [27]. Palmitoleic acid protects against hypertension [28]. |
| Lactic acid (Liver) | In animals, L-lactate is constantly produced from pyruvate via the enzyme lactate dehydrogenase in a process of fermentation during normal metabolism and exercise. Exogenous food such as beverages. | Gluconeogenesis; Pyruvate metabolism | Aging (Up) | - |
| Lithocholic acid (Liver) | It is formed from chenodeoxycholate by bacterial action and is usually conjugated with glycine or taurine. | Secondary bile acid; It acts as a detergent to solubilize fats for absorption. | Aging (Down) | Lithocholic acid inhibits inflammation and regulates metabolism [29]. |
| Pyruvate (Serum) | Pyruvate is found in all living organisms ranging from bacteria to plants to humans. It | Urea cycle; Glucose-alanine cycle; | Aging (Up) | - |

| | | | | |
|---|---|---|---|---|
| | is intermediate compound in the metabolism of carbohydrates, proteins, and fats. | Glycine, serine, and threonine metabolism | | |
| 1.3-Dihydroxyacetone (Serum) | It is often derived from plant sources such as sugar beets and sugar cane, by the fermentation of glycerin. | In combination with naphthoquinones, it acts as a sun screening agent. | Aging (Down) | - |
| Acetone (Serum) | Acetone is produced and disposed of in the human body through normal metabolic processes. It is normally present in blood and urine. | Ketone body metabolism | Aging (Down)<br>FXR KO (Down) | - |
| Methylamine (Urine) | Methylamine occurs endogenously from amine catabolism; Methylamines produced by microbial metabolism of dietary choline and L-carnitine.<br>Exogenous food such as teas | Tyrosine metabolism | Aging (Down) | - |
| N.N-Dimethylglycine (Urine) | A derivative of the amino acid glycine | A microbial metabolite; A biomarker for the consumption of legumes | Aging (Down)<br>FXR KO (Up) | - |
| Betaine (Urine) | Exogenous food such as fruits (e.g., berries) | Glycine, serine, and threonine metabolism; Methionine metabolism | Aging (Up) | Betaine inhibits inflammation and apoptosis [30]. |

| | | | | |
|---|---|---|---|---|
| 2-Hydroxyisobutyrate (Urine) | Non-essential, secondary metabolite | May serve as defense or signaling molecules. | Aging (Up)<br>FXR KO (Up) | - |
| Sn-Glycero-3-phosphocholine (Urine) | Formed in the breakdown of phosphatidylcholine | One of the four major organic osmolytes in renal medullary cells, changing their intracellular osmolyte concentration in parallel with extracellular tonicity during cellular osmoadaptation.<br>Retinol metabolism | Aging (Up)<br>FXR KO (Up) | - |
| 3-Indoxylsulfate (Urine) | A metabolite of the common amino acid tryptophan and is derived through the consumption, digestion, and microbial processing of protein-rich foods. Indoxyl sulfate is technically a bacterial co-metabolite, meaning that it is derived from both bacterial and host metabolism. | A uremic toxin and cardiotoxin | Aging (Up) | - |
| Ascorbate Vitamin C (Urine) | An essential nutrient in human diets, including citrus fruits and many vegetables; | A water-soluble vitamin; Antioxidant; An electron donor for enzymes involved in collagen | Aging (Down) | Ascorbate ameliorates factors linked to Alzheimer's disease pathogenesis [31]. |

| | | | | |
|---|---|---|---|---|
| | A microbial metabolite produced by Ketogulonicigenium | hydroxylation, biosynthesis of carnitine and norepinephrine. | | |
| Melibiose (Liver) | This sugar is produced and metabolized only by enteric and lactic acid bacteria and other microbes, such as *Escherichia*. It is not an endogenous metabolite but may be obtained from the consumption of partially fermented molasses, brown sugar, or honey. | Galactose metabolism | Aging (Up), FXR KO (Down) | - |
| Glucoheptulose (Liver) | L-arabinose | - | FXR KO (Down) | - |
| UDP-N-acetylglucosamine (Liver) | Exogenous, food such as animal food | Glucose sensor; Amino sugar metabolism; Elevated UDP-N-acetylglucosamine has an effect on insulin-stimulated glucose uptake. | FXR KO (Up) | - |
| Uridine (Liver) | Synthesized from uracil; Uridine is found in many foods (anything containing RNA). | Pyrimidine metabolism | FXR KO (Up) | Uridine alleviates carbon tetrachloride induced liver fibrosis [24]. |

| | | | | |
|---|---|---|---|---|
| Isomaltose (Liver) | A product of the caramelization of glucose; Exogenous, food such as beverages | Starch and sucrose metabolism; Metabolic pathways | FXR KO (Up) | Isomaltulose improves insulin resistance in NAFLD patients [32]. |
| Ribose-5-phosphate (Liver) | Exogenous, food such as fruits; A product and an intermediate of the pentose phosphate pathway. | Purine metabolism; Pentose phosphate pathway | FXR KO (Down) | Ribose-5-phosphate isomerase A overexpression induces oncogenesis in HCC [33]. |
| 2-hydroxybutanoic acid (Liver) | Primarily produced in mammalian hepatic tissues that catabolize L-threonine or synthesize glutathione. Exogenous, food such as animal foods | An early marker for both insulin resistance and impaired glucose regulation | FXR KO (Down) | - |
| Arachidic acid (Liver) | A minor constituent of butter, perilla oil, peanut oil, corn oil, and cocoa butter. | Saturated, long-chain fatty acids | FXR KO (Down) | Arachidic acid induces liver dysfunction in hyperglycaemic rats [34]. |
| Malic acid (Liver, serum) | Exogenous, food such as herbs and spices | Citric acid cycle; Gluconeogenesis; Pyruvate metabolism; Malate-aspartate shuttle | FXR KO (Up in the liver) FXR KO (Down in the serum) | Malic acid is increased in NAFLD [35] |
| Succinate (Serum, Urine) | Exogenous, food such as fruits | A cell signaling molecule; Succinate alters gene expression patterns, thereby modulating the epigenetic landscape or it | FXR KO (Down in serum and up in urine) | Elevated extracellular succinate in liver tissue drives inflammation [36]. |

| | | | | |
|---|---|---|---|---|
| | | can exhibit hormone-like signaling functions. | | |
| Alanine (Serum) | It is formed in vivo by the degradation of dihydrouracil and carnosine. | A neurotoxin; A mitochondrial toxin; A metabotoxin; Beta-alanine metabolism | FXR KO (Down) | - |
| Glutamine (Serum) | Non-essential amino acid | Proteinogenic $\alpha$-amino acid; Glutamate metabolism; Purine metabolism; Urea cycle | FXR KO (Up) | Glutamine is beneficial for human inflammatory bowel disease [37]. |
| Phenylalanine (Serum) | An essential amino acid and the precursor of the amino acid tyrosine, highly concentrated in high protein foods, such as meat, cottage cheese, and wheat germ. An additional dietary source of phenylalanine is artificial sweeteners containing aspartame. | Proteinogenic $\alpha$-amino acid; A precursor for catecholamines including tyramine, dopamine, epinephrine, and norepinephrine; A neurotoxin and a metabotoxin. | FXR KO (Up) | - |
| Tyrosine (Serum) | Exogenous, food such as fruits | Proteinogenic $\alpha$-amino acid; Tyrosine metabolism | FXR KO (Down) | - |
| Glucose (Serum) | Exogenous, food such as fruits | Primary source of energy for all living organisms, Glycolysis; Gluconeogenesis; Lactose synthesis | FXR KO (Up) | - |

| | | | | |
|---|---|---|---|---|
| Creatinine (Urine) | Exogenous, food such as herbs and spices | An amino acid derivative; A waste product and is normally eliminated in large quantities by the kidneys through urinary excretion. | FXR KO (Down) | - |
| Taurine (Urine) | An essential amino acid; Foods such as vegetables, animal and fish protein | A neurotransmitter in the brain; Taurine and hypotaurine metabolism; Primary bile acid biosynthesis | FXR KO (Down) | Taurine displays potential ameliorating effects against different neurological disorders [38]. |
| N-Phenylacetylglycine (Urine) | Exogenous, food such as animal food | A surrogate biomarker for phospholipidosis | FXR KO (Up) | A biomarker for dimethylnitrosamine-induced hepatic fibrosis in a rat model [39]. |
| Guanidoacetate (Urine) | Exists naturally in all vertebrates; Formed primarily in the kidneys by transferring the guanidine. | $\alpha$-amino acid and derivatives; Arginine and proline metabolism; Glycine, serine, and threonine metabolism | FXR KO (Down) | - |
| trans-Aconitate (Urine) | Normally present in human urine; Detected in foods, such as garden tomato fruits, root vegetables, soybeans, and rice. | A biomarker for the consumption of soy products | FXR KO (Down) | - |
| Cis-Aconitate (Urine) | Cow milk | Glutaminolysis and cancer pathway; Citric acid cycle | FXR KO (Down) | - |

| | | | | |
|---|---|---|---|---|
| N.N-Dimethylglycine (Urine) | Exogenous, food such as animal foods; An amino acid derivative found in the cells of all plants and animals and can be obtained in the diet in small amounts from grains and meat; A byproduct of homocysteine metabolism; A microbial metabolite | A biomarker for the consumption of legumes; Glycine, serine, and threonine metabolism; Methionine metabolism | FXR KO (Up) | Plasma N.N-Dimethylglycine is markedly decreased in Alzheimer's disease patients compared with normal controls [40]. |

HCC, hepatocellular carcinoma; NAFLD, nonalcoholic liver disease; -, unknown
